# On the Effectiveness of UAS for Anti-Poaching in the African Arid Savanna

**DOI:** 10.1101/660126

**Authors:** Marlice van Vuuren, Rudie van Vuuren, Graham D. Lutz, Larry M. Silverberg

## Abstract

This paper describes a field study that examined the effectiveness of unmanned aerial vehicles (UAV) in anti-poaching enforcement in parks and game reserves. In the field study, a UAV attempted to spot mock poachers while the mock poachers tried to spot the UAV. The field study was conducted at N/a’an ku sê, an operational game reserve in the central region of Namibia. In total, 118 trials were completed, providing 236 UAV-poacher interdiction scenarios. Of these, 198 were during the day, 152 with a quadcopter and 46 with a fixed-wing. Live spotting success during the day varied due to the hiding behavior of the mock poachers, with the highest and lowest success rates of spotting being 86% for poachers in the open and 25% for poachers hiding under canopy cover. The UAVs were demonstrated to be a potentially effective tool for anti-poaching patrol and interdiction, in part, because of their ability to spot poachers. The pursuit of integrating the UAV into current anti-poaching patrol and interdiction efforts in arid savanna landscapes is strongly recommended.

## Introduction

The high rate of animal extinction of such high-value African species, such as rhinoceros and elephant, is credited to poaching (Western 1985; Douglas 1987). They are illegally hunted for their horns and tusks, and sold for the production of traditional medicines and for items of cultural status. From 1990 to the present, the African elephant population decreased by 90% and between 1960 and 1990 the black rhino population decreased by 95% (Kamminga, 2018). To combat the decline of high-value African species, effective anti-poaching policies require strong support by local communities, strong anti-poaching efforts by law enforcement, and strong prosecution by legal systems. In law enforcement, the focus is on best practices in patrol and interdiction. Today’s tools include the camera trap, ground surveillance (walking and driving), and aircraft surveillance with the latter often being unaffordable. The advent of the unmanned aerial vehicle (UAV) offers law enforcement with a new, potentially powerful tool.

To date, published research on the effectiveness of the UAV in anti-poaching enforcement is scant. Exceptions include anti-poaching UAV research on 13 farms across South Africa with 20 total trials (Mulero-Pazmany, 2014). Their research focused on image quality of three types of cameras mounted on a fixed-wing UAV for day and nighttime trials. They found some viability in using UAVs as an anti-poaching tool to spot both rhinos and humans. Also, a security group implemented an anti-poaching UAV program using a fixed-wing UAV as a nighttime anti-poaching tool (Air Shepherd, 2018). They reported statistics regarding their successes, noting that in one area in which 19 rhinoceros were once killed, there were no killings after deploying their UAVs. Information on their methodology, however, was not released. Ultimately, the role of the UAV in enforcement, despite early signs of great promise, is not yet settled. How will the UAV compliment camera traps, walking, and driving at parks and game reserves? What overall level of training and effort will be required? At the root of these questions is the more basic question, which this paper examines, of the relative abilities of the UAVs and the poachers to sense, by sight and sound, each other’s presence.

The method section describes a field study in which a UAV attempts to spot mock poachers while the mock poachers try to spot the UAV. The different factors that impact the ability to spot one another are discussed in the results section and the implications of the study to patrol and interdiction are described in the summary and conclusions section.

## Method

The objective of this study was to examine the anti-poaching UAV and the poacher’s abilities to sense, by sight and sound, each other’s presence. These abilities, once understood, could be applied to the patrol scenario, where the focus is on protecting an area, and to the interdiction scenario, where the focus is on apprehending poachers. Indeed, the parameters that characterize the abilities and constraints of sensing each other’s presence will drive how to best integrate the UAV into patrol and interdiction. Currently, the three most popular methods employ (1) camera traps, (2) walking, and (3) driving. Today, camera traps are most commonly used as a way to collect information after-the-fact, to use in prosecution and to learn poacher habits. Such countries as Tanzania and Borneo have used them effectively. Collecting real-time data with them, on the other hand, has been difficult due to technical hurdles and manpower requirements to process data. Also, camera traps must be hidden from the view of the poachers and covering large, open savannas with camera traps is problematic. Walking and driving are currently the two most popular methods of patrolling and interdicting poachers. A pair of well-trained security personnel can walk 3 to 5 km in a day without being detected. The areas in which they walk are unrestricted by roads and they can exploit all of their senses. On the other hand, security personnel can cover much less area on foot than by ground vehicle and walking is particularly dangerous when confronting a poacher. In a ground vehicle, a pair of security personnel routinely covers 75 km in a day. In contrast with walking, the ground vehicle is readily detected by the poacher so the ground vehicle serves as a deterrent and is most commonly deployed around perimeters to areas. In interdiction, it serves as a rapid means of getting close to an area and as a means to herd poachers to desirable locations. In light of these considerations, the UAV has the potential of complimenting the walking and driving approaches. In patrol, it can cover an area relatively fast. Before needing its battery changed, a simple, low-weight radio controlled or autonomous multi-rotor UAV can cover about 8 km in twenty minutes and a comparable fixed-wing UAV can cover 15 km to 20 km in 30 minutes. Like the ground vehicle, the UAV collects visual data but, unlike walking, it does not capitalize on the other senses. In patrol, the UAV is well-suited to following perimeters. It can cover a much wider swath than the ground vehicle can and its vantage point from the air is much better than the vantage point from the ground. As a result, the UAV can protect a perimeter much better than a ground vehicle can. In interdiction, it can assist walking and ground vehicle personnel by providing them with important information about the poachers, such as their number, arms, and heading. Since the UAV is remote, it also brings greater safety to both patrol and interdiction.

The basic question is therefore this: What are the practical constraints of the UAV with regard to the comparative abilities of the UAV and the poacher to sense each other’s presence? These abilities, once understood, could then be applied to the patrol scenario, where the focus is on protecting an area, and could be applied to the interdiction scenario, where the focus is on apprehending poachers. Note that this paper provides general conclusions about the effectiveness of the UAV in the patrol and interdiction scenarios. Specific scenarios are laid out depending on such factors as terrain, existing roads, and complimentary resources available, so detailed conclusions and recommendations are beyond the scope of this paper.

Fig 1 below depicts the test area located at N/a’an ku sê, an operational game reserve in the central region of Namibia.

**Figure 1:**
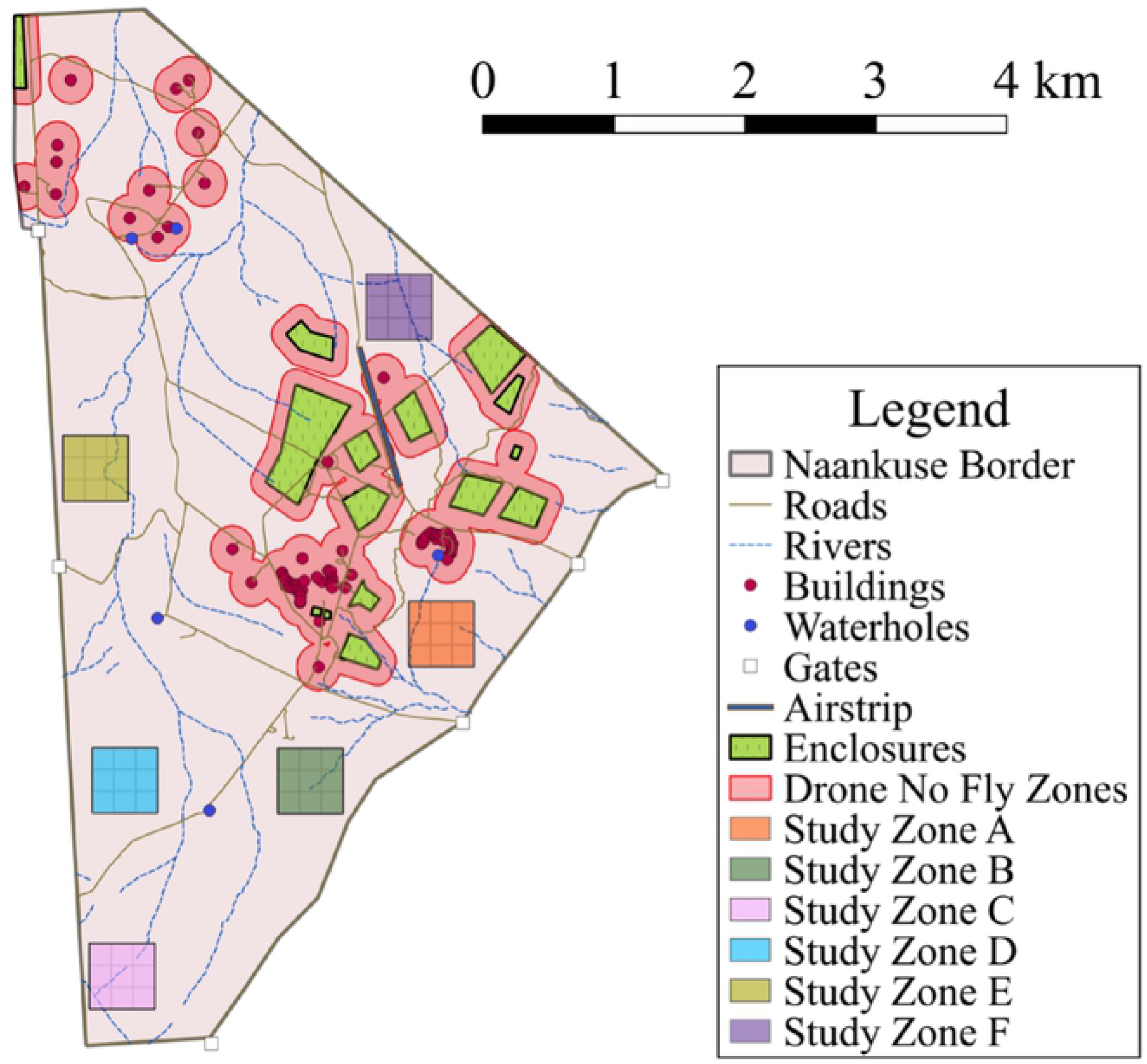
N/a’an ku sê Map and Study Zones

As shown, the test area was divided into six study zones. Each had differences in terrain and vegetation cover. Each measured 500m by 500m which was sufficiently large to challenge the UAVs and the mock poachers and which was sufficiently small to enable the sighting to be accomplished in an appropriately small amount of time. As shown, each of the zones was further subdivided into a grid of nine sectors, each sector serving as a hiding area for the mock poachers. Three hiding behaviors were defined as follows:

1. In the open: *uncovered by and separated from any vegetation by at least 5 meters*
2. Amongst the bush*: within five meters or less of vegetation but with a clear view of the sky above*
3. Under the canopy: *completely underneath the cover of bush*

The tests were performed over a 10-week period in 2018. Each test was accomplished by a six-person field unit consisting of two mock poachers, a pilot, a co-pilot, a note-taker and an observer. The roles were randomized after each trial to reduce biases resulting from variations in skill level. The pilot was responsible for setting up the radio control equipment, launching and landing the UAV, and video observation. The co-pilot was responsible for the initial UAV setup and video observation. The note-taker was responsible for recording all data, before, during and after the test. The observer maintained situational awareness and visual contact with the UAV and operated the radio for safety purposes. In each test, the mock poachers were assigned to a random sector and to a random hiding behavior.

Before the beginning of each test, role assignments, wind speed and direction, cloud cover, flight altitude, flight speed, UAV camera angle, and GPS takeoff location were recorded. Next, at the beginning of each test, the field unit set up its equipment beside the takeoff location while the mock poachers went to their assigned random hiding sectors and assumed their hiding behaviors. Then, upon takeoff, the note taker started a stopwatch and the observer told the poachers via radio to start their stopwatch. The two synced stopwatches were later used to correlate UAV and mock poacher observations. After takeoff, the UAV autonomously flew a pre-programmed search pattern, with the pilot and co-pilot continuously monitoring a live video feed. Any time the pilot and co-pilot thought they spotted a mock poacher, the time was recorded. During the test, the mock poachers recorded the GPS coordinates of their hiding spot and their assigned hiding behavior and the times they first heard the UAV, saw the UAV, and when the UAV was directly overhead. The note-taker entered this data into the dataset at the conclusion of the test.

A post processing review of photos and recorded video allowed the sightings to be either confirmed or marked as false and it enabled sightings that the pilot and the co-pilot missed to be identified.

Three UAV systems were chosen: two quadcopter systems and a fixed-wing UAV system. They were meant to be representative of the different systems that would be employed in anti-poaching efforts. Table 1 describes the technical features of the UAV systems.

**Table 1:**
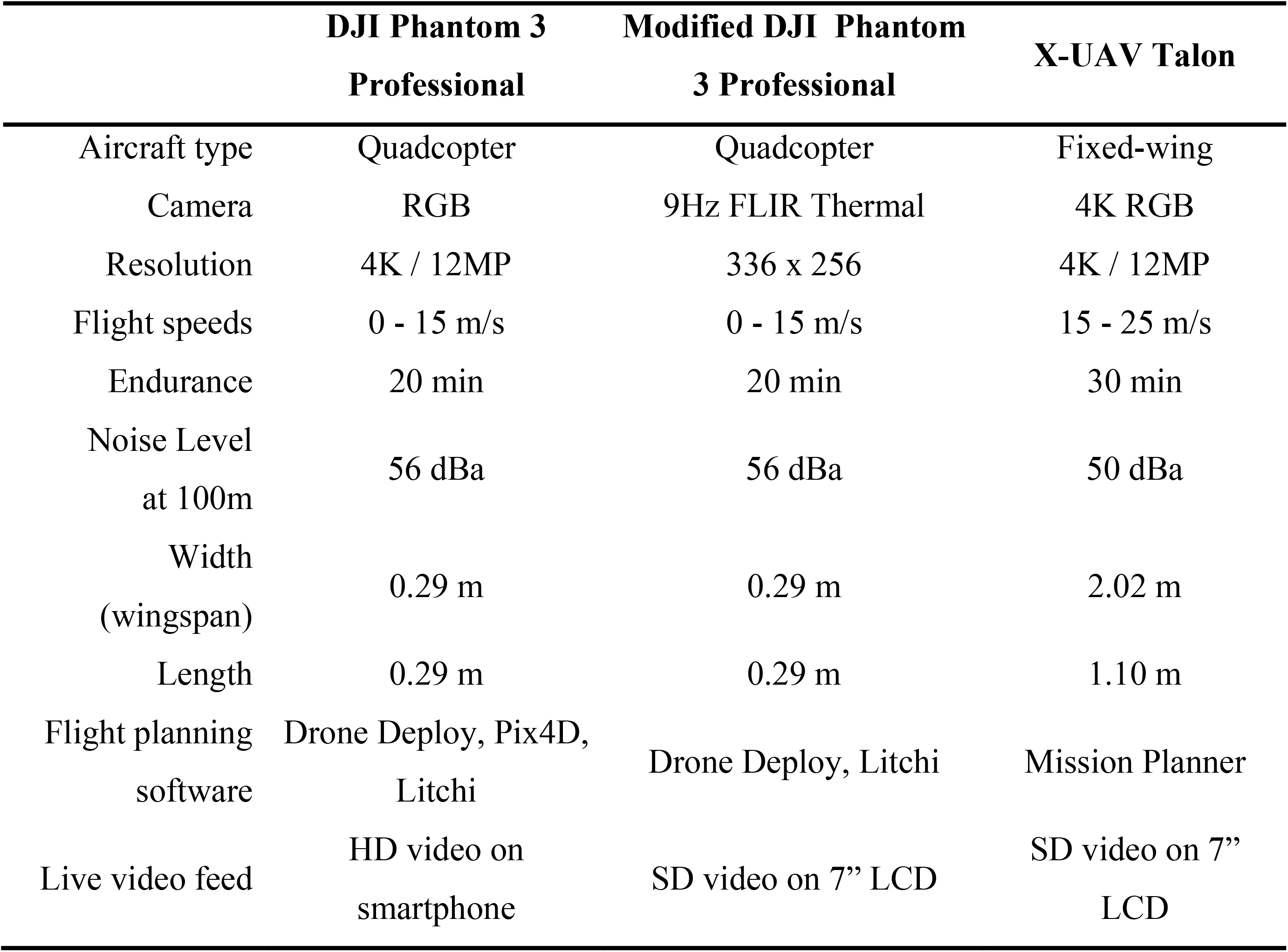
UAV Technical Descriptions

The parallel track search method was employed during the tests (WSDOT, 2008). The parallel tracks were flown in the north-south directions to prevent flying into and away from the sun. The ability to adjust parameters throughout the flights was also necessary to obtain the best results. The adjusted parameters were camera angle, flight speed, altitude, and recording mode (stills and video). For the fixed-wing system, a modified version of the parallel track search method was flown to account for its limited turning radius. The difference in flight paths is shown in Fig. 2 below.

**Figure 2:**
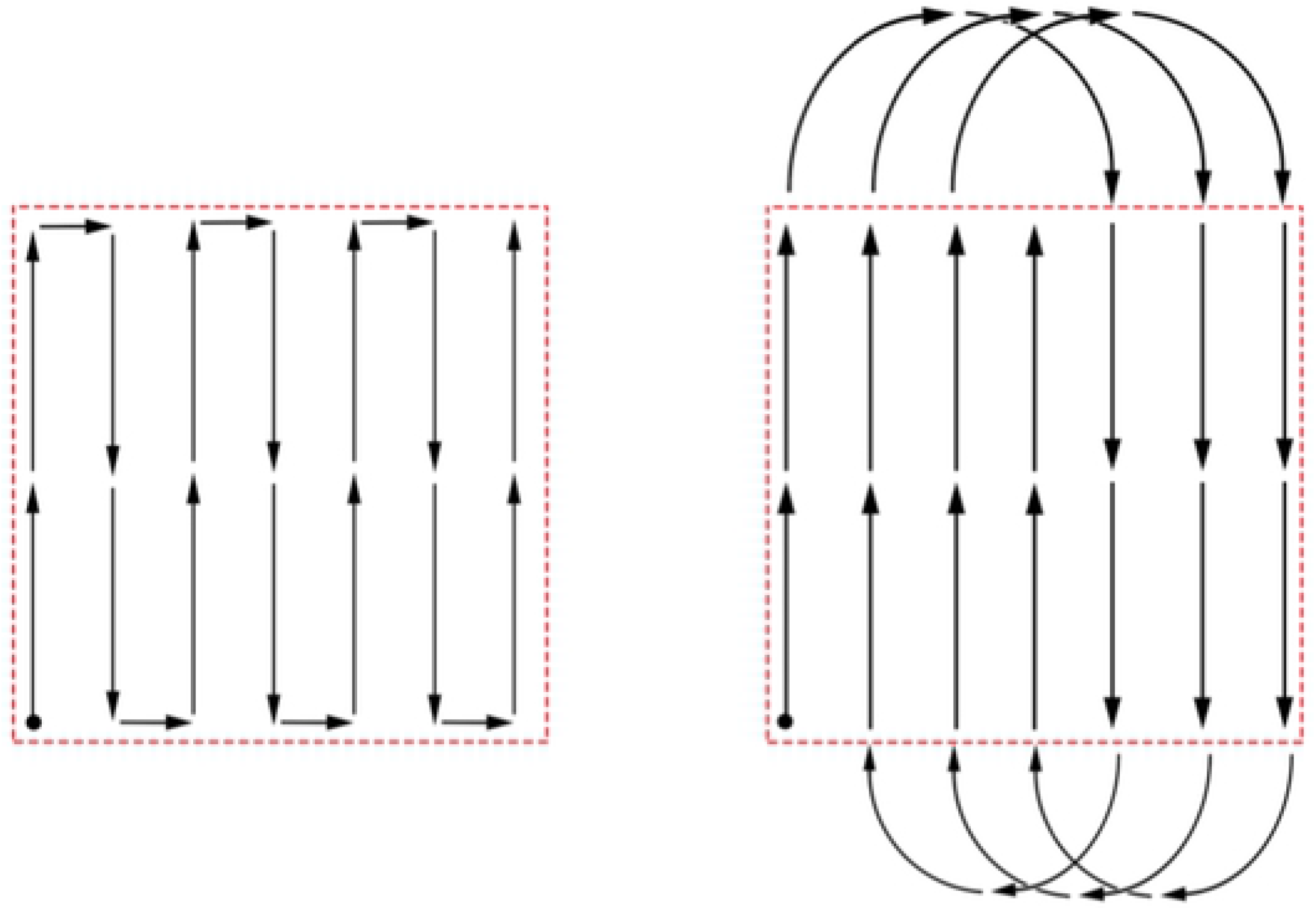
Parallel track search method for the quadrotors (left) and its modification for the fixed-wing UAV (right)

The quadcopter systems required two batteries to complete the search pattern, returning to the takeoff location to change batteries half way through the trial. The fixed-wing system was capable of completing two search patterns per battery, and, to minimize wear and tear on the UAV, was allowed to circle, rather than land between tests. In the night tests using the quadcopter system with a FLIR camera, the onboard lights were blacked out except for the navigation lights, which were allowed to remain turned on during takeoff and landing. Several color pallets were available for the thermal imagery however, after testing, the “white hot” pallet was chosen because it provided the most definition; the humans stood out best with it.

An even number of tests were conducted in each of the 6 zones so that the environmental factors, such as vegetation cover and terrain, would be evenly represented. Daytime flights were conducted between the hours of 9:00 and 17:00. The thermal camera required ground temperatures that are cool relative to the body temperature so the flights with the thermal camera were conducted at least 30 minutes or more after dusk.

## Results

In total, 118 trials were completed with 236 mock poachers in the field. From the trails, information was gleaned about the abilities of the UAVs to spot poachers in the three different hiding behaviors, and the comparative abilities of the mock poachers to be the first to spot the UAVs.

Figures 3, 4, and 5 below show illustrative images of mock poachers in the field. In each figure there are 2 poachers. In Figs. 3 and 4, one is in the open and the other is amongst the bush. In Fig. 5, one is amongst the bush and the other is under a canopy. In all of the tests, the camera angles were measured from nadir, with any angles deviating from nadir in the forward facing direction of the UAS. Figures 3 and 4 compare the nadir and 45° off nadir camera angles, respectively, showing that the 45° camera angle is preferred over the nadir camera angle due to the increased visibility of the human.

**Figure 3:**
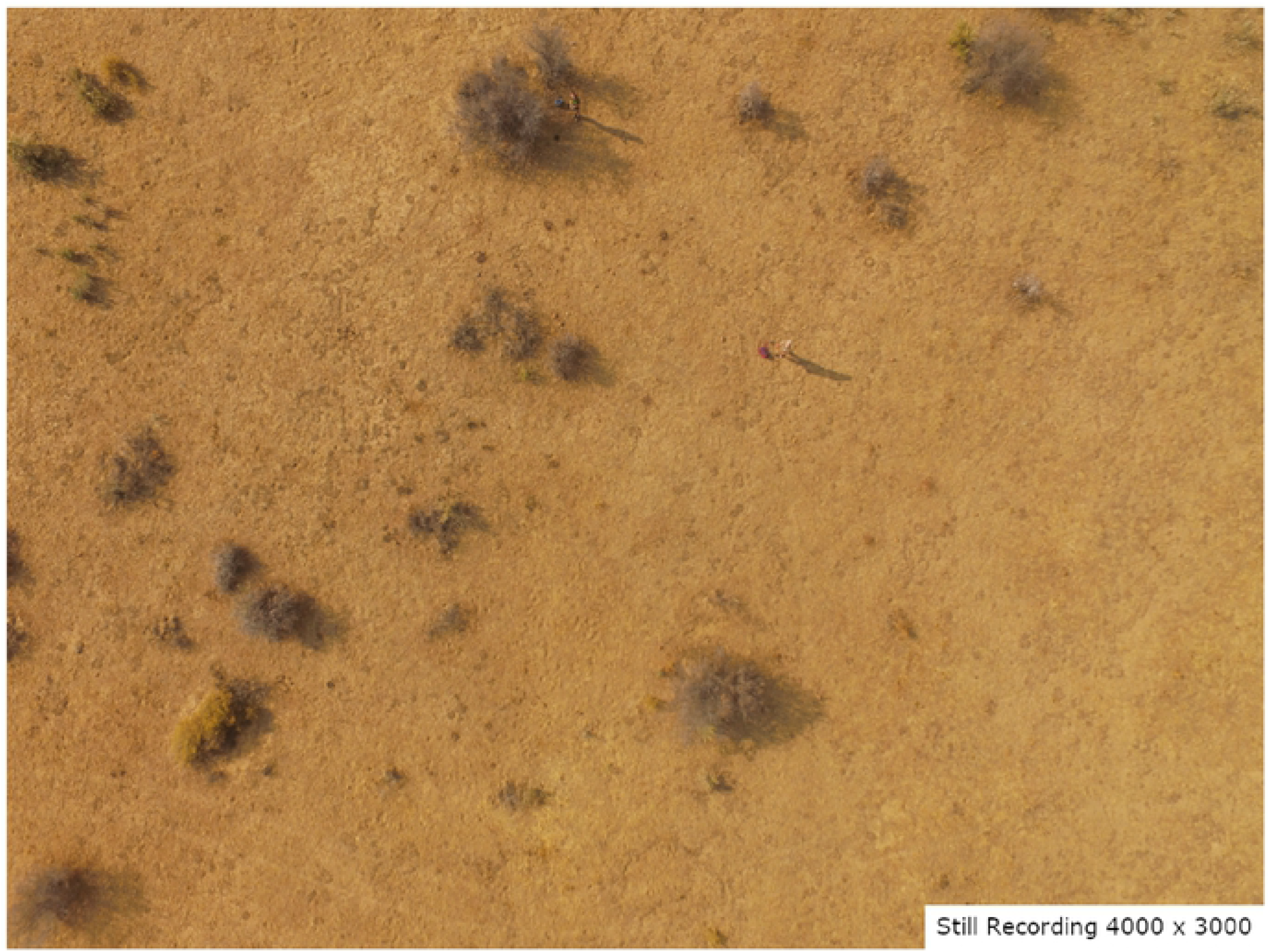
Mock poachers, 0 degree nadir camera angle

**Figure 4:**
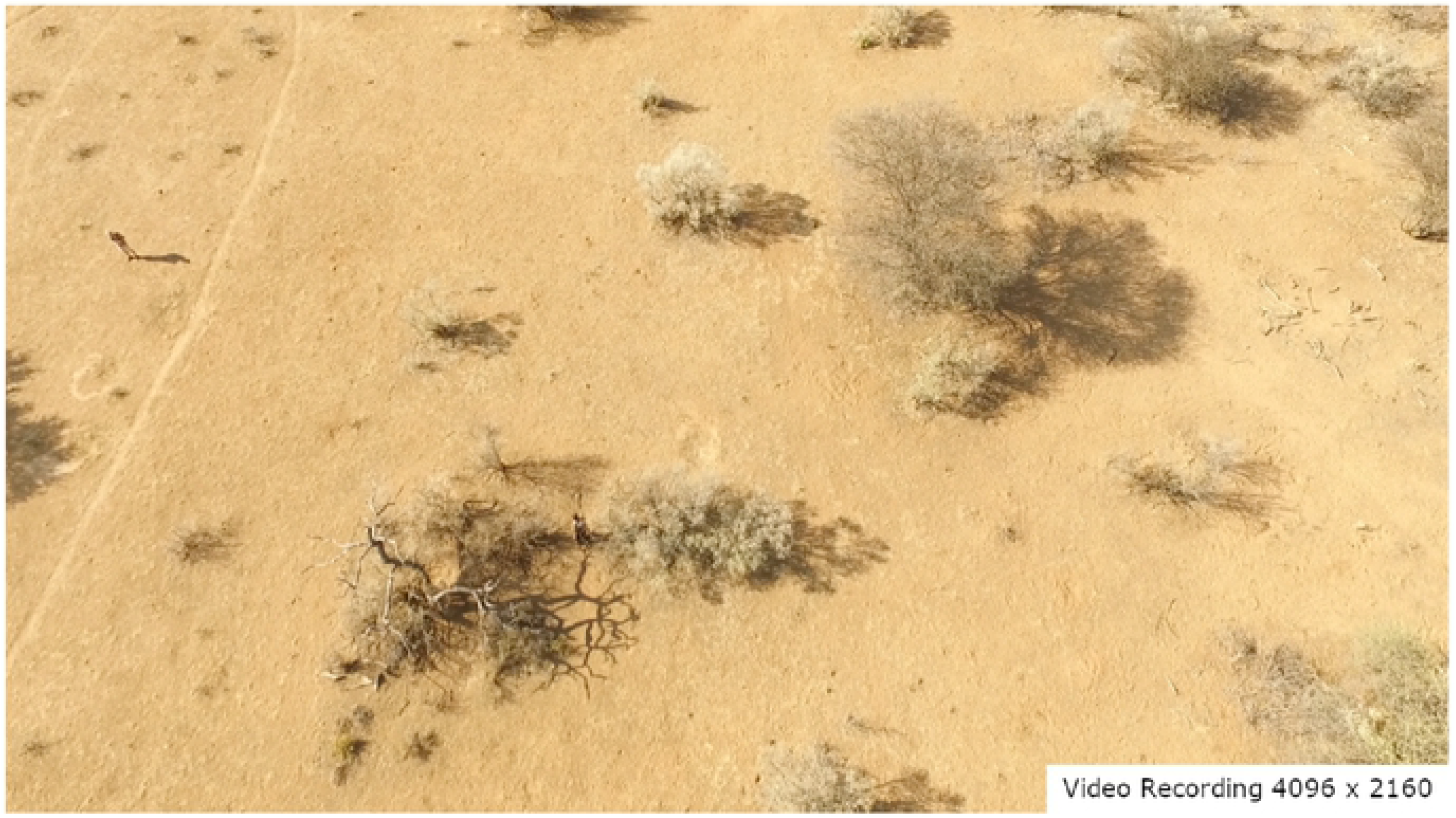
Mock poachers, 45 degree off nadir camera angle

**Figure 5:**
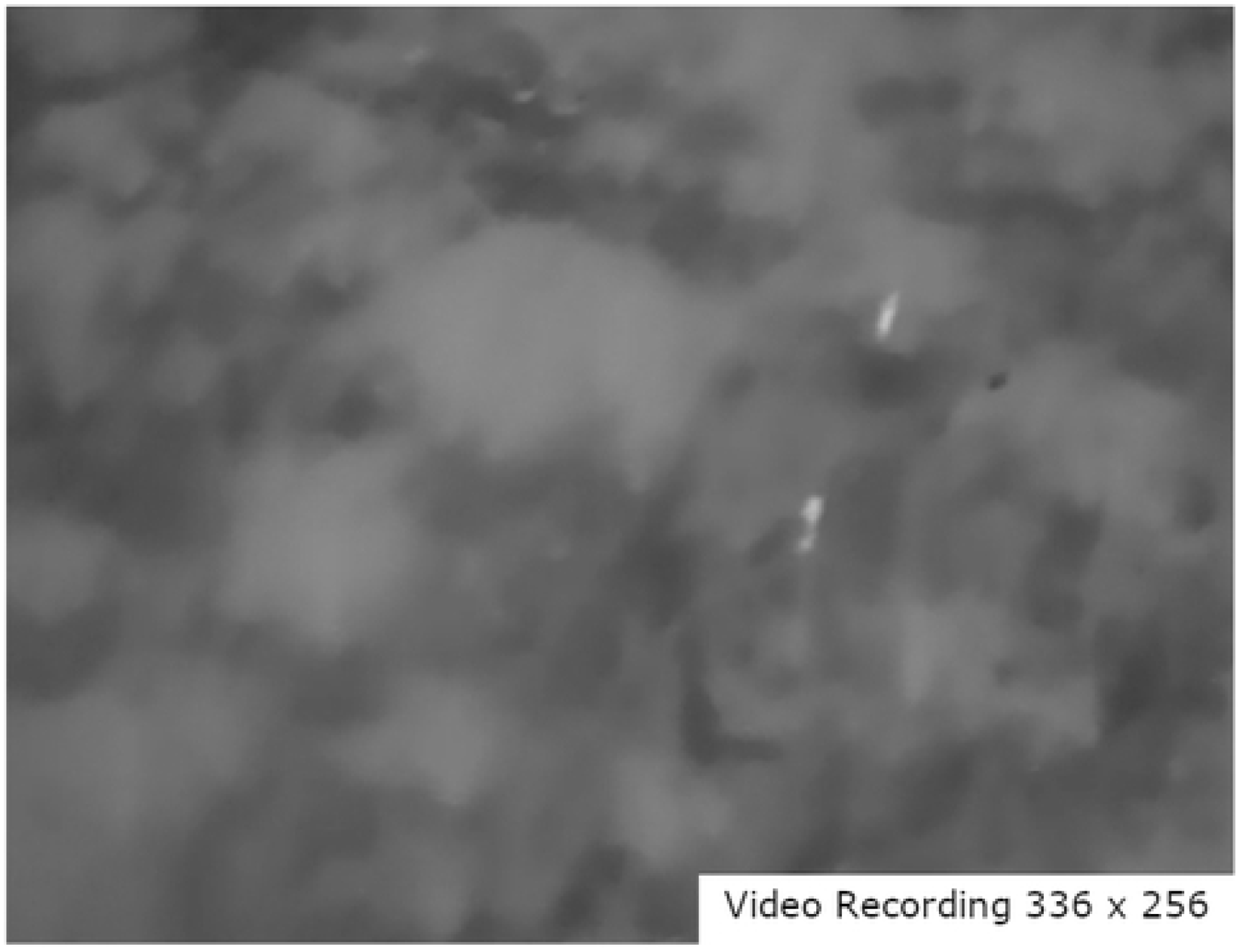
Mock poachers, FLIR camera, 45 degree off nadir camera angle

For still pictures, camera angles of both nadir and 30° off nadir were considered. For flights in which video was recorded, camera angles of 30° off nadir and 45° off nadir were used. Table 2 below tabulates the successes rates for the different camera angles and recording modes. Video recording and the 45° off nadir camera angles yielded the best results and were also preferred by all of the observers.

**Table 2:**
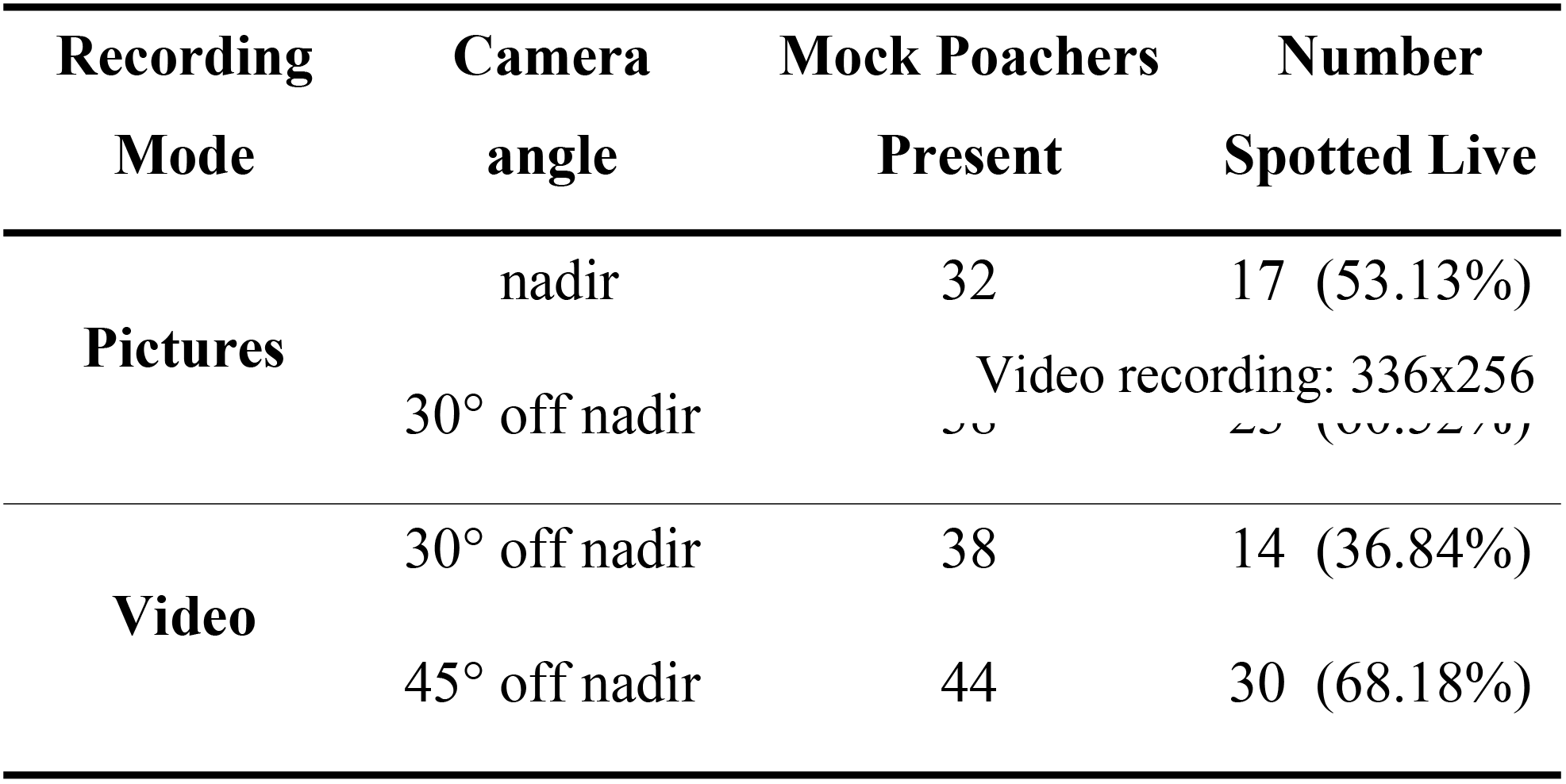
Spotting success with different camera angles and recording types

Observers preferred the video recording to pictures because the picture mode was distracting; live video feed flashed black every time a picture was taken. For camera angles, 45° off nadir was preferred because it allowed the human’s profile to be seen which allowed for better identification than a 0° nadir view. The 45° off nadir angle gave the observers a better chance of seeing under the canopy and also resulted in mock poachers being on screen for a longer time than the 30° off nadir or 0° nadir camera angles. In their work, Mulero-Pazmany, et al. tested camera angles of 15° off nadir and 60° off nadir, however no mention was made as to which was superior.

Flight speeds for all of the trials ranged from 4.7 m/s to 9.5 m/s, as shown in Table 3. These were divided into three ranges with the majority of the trials falling into the lower two of the three ranges. This is because live observers immediately noticed that in the fastest range the UAV was moving too fast for them to effectively scan the live video feed for poachers. Despite the percent of mock poachers spotted being almost identical for the two slower flight speeds, all of the observers found that the slower flight speed made their job feel much easier and less stressful.

**Table 3:** Spotting success with different flight speeds

| **Flight**<br>**speed** | **Mock Poachers**<br>**Present** | **Number**<br>**Spotted Live** |
| --- | --- | --- |
| **4.7-6 m/s** | 64 | 36 (56.25%) |
| **6.1-7.5 m/s** | 78 | 44 (56.41%) |
| **7.6-9.5 m/s** | 10 | 4 (40.00%) |

Table 4 below shows the success rates for spotting poachers in the field during the daytime using both the fixed-wing and quad platforms. Overall the quadcopter system was more successful than the fixed-wing system with both systems having the highest success spotting poachers in the open and the least success spotting poachers under the canopy. The hiding behavior had a very strong effect on the ability of the UAV system and the live observer to spot the mock poacher, as expected, with the mock poachers under the canopy being 2 to 3 times as hard to spot as those in the open.

**Table 4:** Spotting success by hiding behavior for fixed-wing and quadcopter

| **Hiding**<br>**Behavior** | **Mock Poachers Present** |  | **Number Spotted Live** |  |
| --- | --- | --- | --- | --- |
|  | Quadcopter | Fixed-wing | Quadcopter | Fixed-wing |
| **Open** | 44 | 13 | 38 (86.36%) | 9 (69.23%) |
| **Bush** | 44 | 17 | 30 (68.18%) | 7 (41.18%) |
| **Canopy** | 64 | 16 | 16 (25.00%) | 5 (31.25%) |

The observers watching the live video noted that spotting with the fixed-wing was more difficult because the camera vibrated; it was not gimbaled to the vehicle. Observers also noted the higher flight speeds of the fixed-wing resulted in their job being both more difficult and tiring. The un-gimbaled camera and the higher flight speeds likely contributed to the lower overall success rates of the fixed-wing platform. It is also noteworthy that the fixed-wing had fewer trials than the quadcopter.

A post processing review of the recorded photos and videos determined whether each of the potential sightings noted by the observers was correct. The post processing review was performed with the knowledge of the time at which the UAV passed over the poacher. The review identified instances in which a mock poacher is visible while the observer fails to see them. Figure 6 compares the success rates for the live spotting mock of poachers in each hiding behavior with the spotting in the post processing review. As shown, not all of the poachers were visible, even in the post processing review and the live observers did not catch all of the poachers that were visible to the UAV.

**Figure 6:**
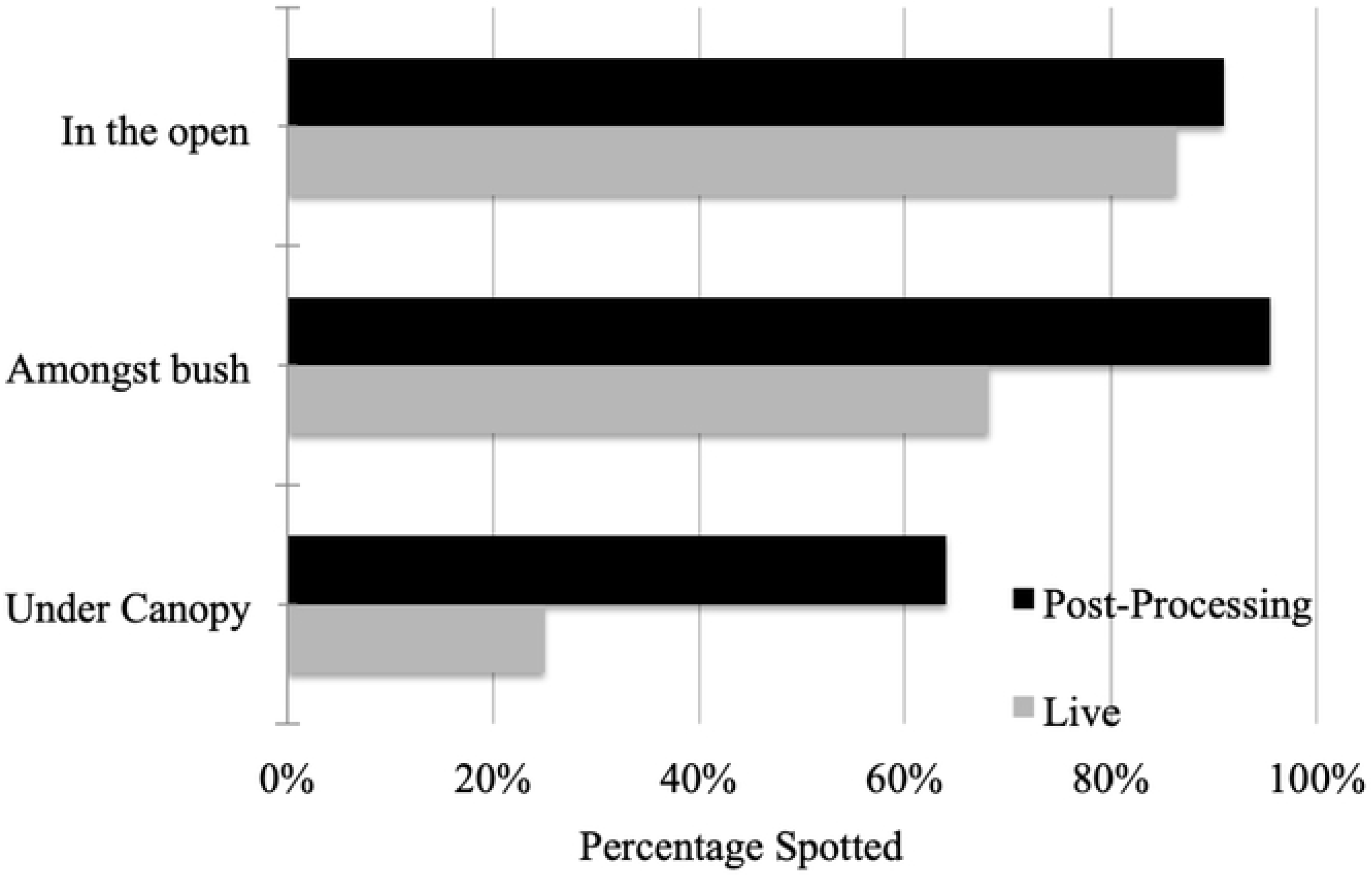
Spotting Live vs. Post-Processing; Quad Daytime

The largest discrepancies between live spotting and post processing occurred when the poachers were fully obscured by the canopy, during which they were spotted in post processing with careful playback. Figure 7 below shows the same live versus post processing comparison as above except it is with the fixed-wing UAV. The fixed-wing UAV shows trends similar to those of the quadcopter UAV, however, with overall lower success rates.

**Figure 7:**
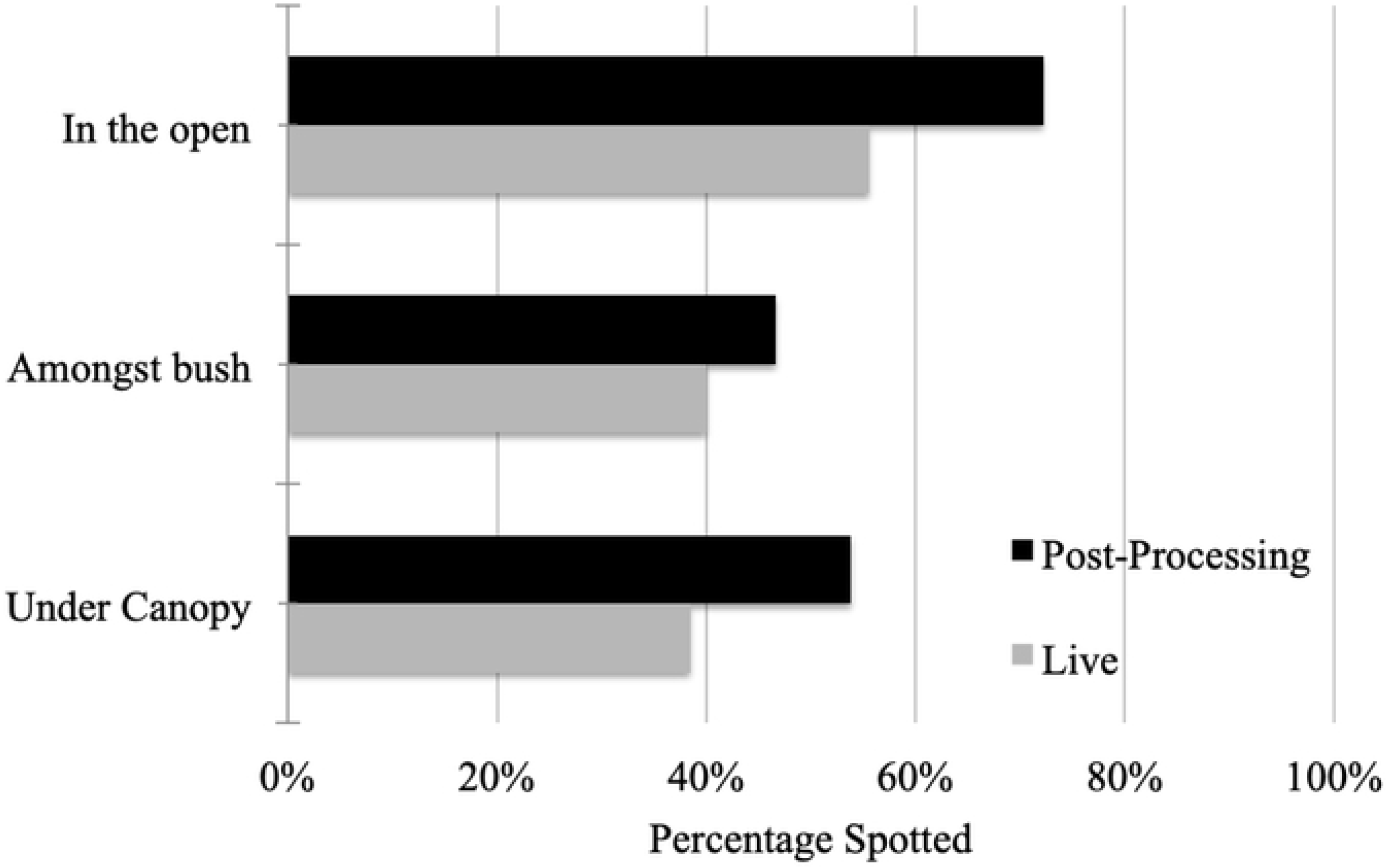
Spotting Live vs. Post-Processing; Fixed-wing Daytime

Figure 8 shows the success rates for spotting poachers using the FLIR camera at night in comparison to those visible in the post processing review. The quadcopter system at night had higher success rates than during the day, spotting all mock poachers in the open and no less than 67% in any of the 3 hiding behaviors.

**Figure 8:**
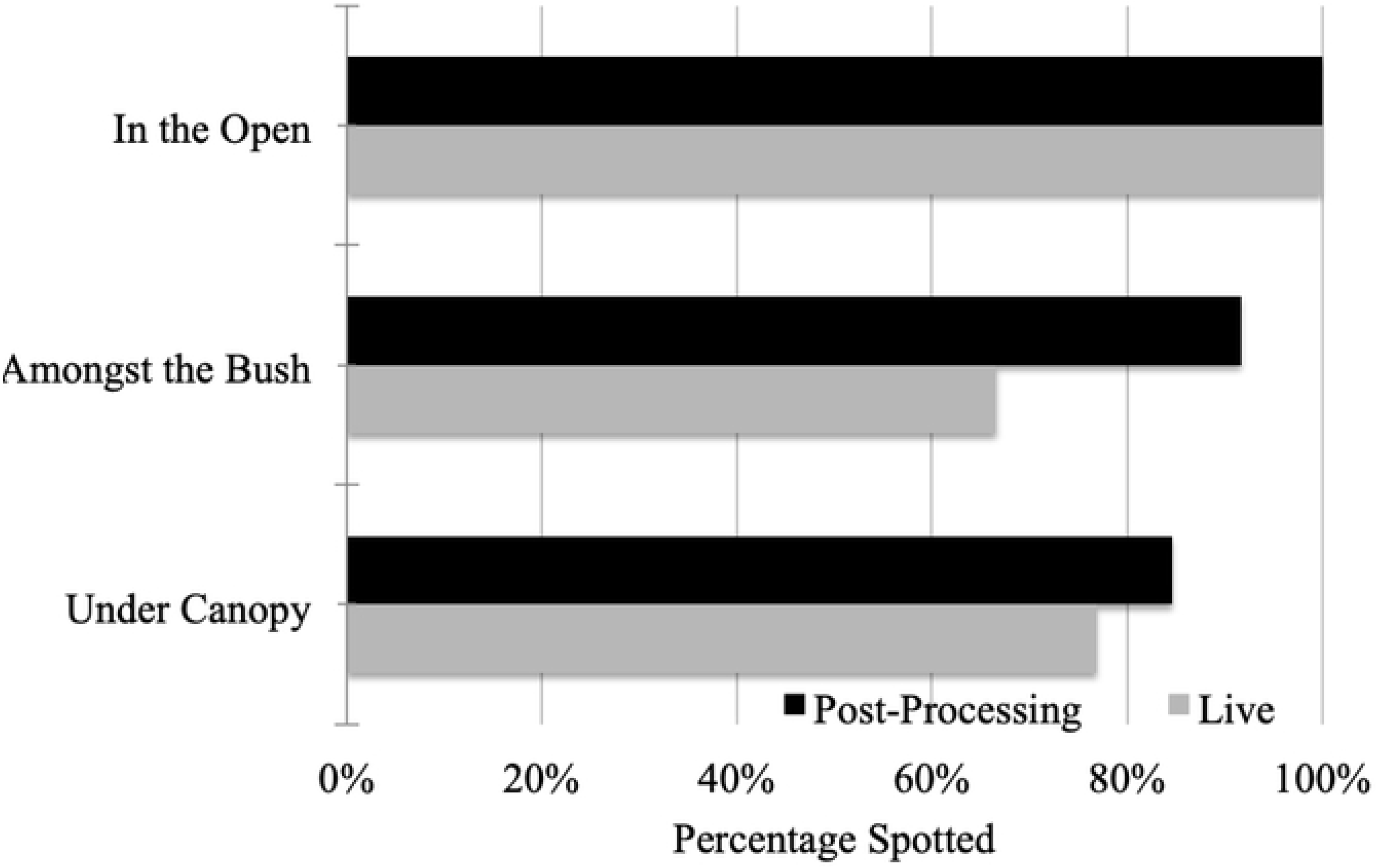
Live Spotting vs. Post-Processing; Quad Night

The high success rates during the night came from the high contrast between human body temperatures and the background. Additionally, the thermal camera almost completely penetrated the canopy unlike the RGB cameras during the day, again allowing for clearer spotting.

In order to assess the reaction time that a poacher or the anti-poaching team would have before the other party was alerted, all of the trials were divided into five categories. During the day, the mock poachers always had the advantage while during the nighttime the UAV, and thus the anti-poaching unit, had the advantage in the majority of the trials. This can be attributed to the blacked out UAV being very difficult to spot. Mock poachers reported that they were really only able to spot the UAV when it flew overhead and when it momentarily passed in front of stars or the moon (See Table 5).

**Table 5:**
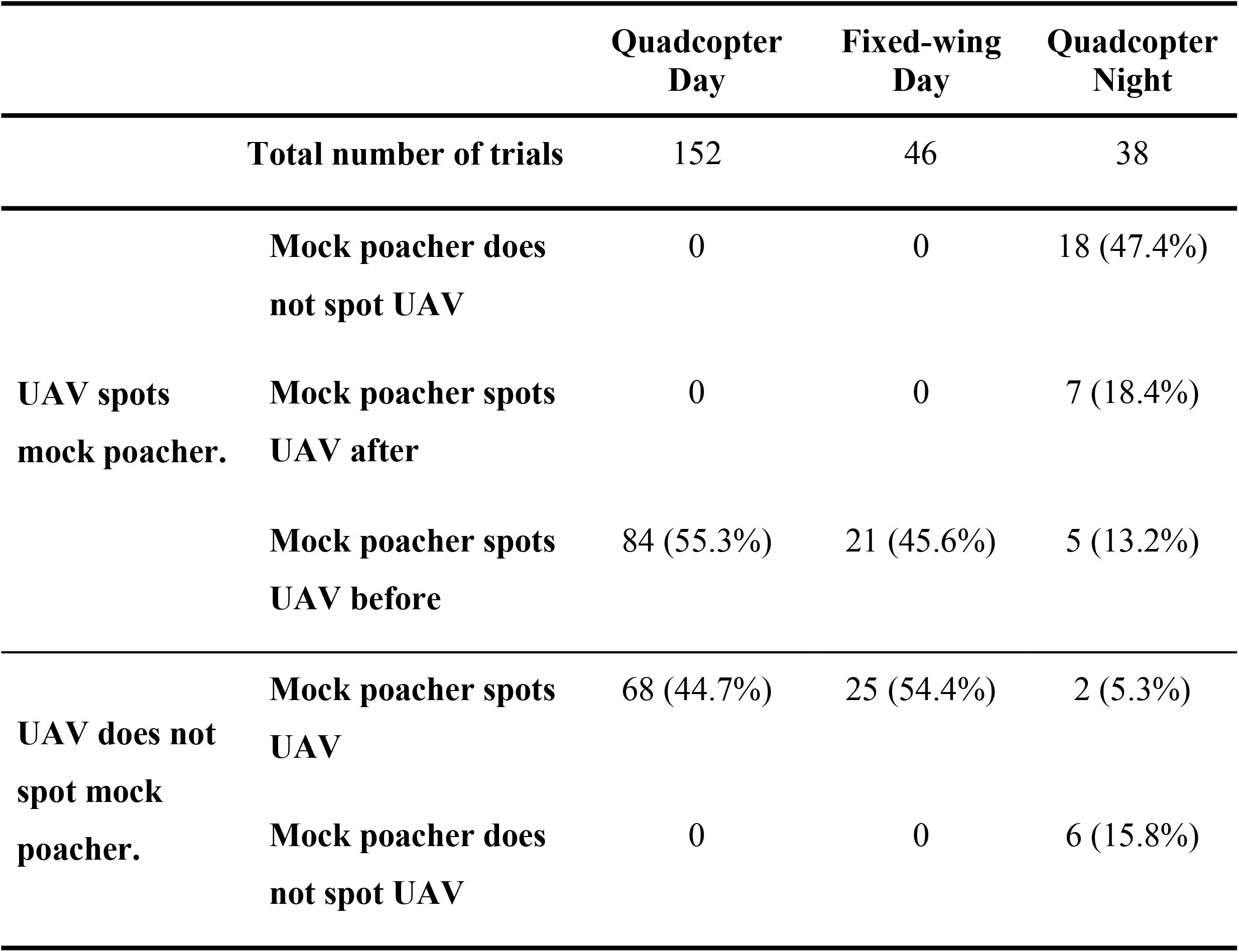
Reaction advantage

For trials in which both the poachers and the UAV spotted each other, the average time difference was calculated. In the day, the average reaction time was 732s in favor of the poachers with a standard deviation of 578s. In the night trials, the average reaction time was 29.3s in favor of the poacher with a standard deviation of 137.7s. The very large standard deviations can be attributed to where the poacher was in relation to the start of the flight pattern since the UAV only sees a poacher as it passes overhead, but the poacher has the advantage of being able to see the UAV whether it is approaching, departing, or passing by laterally.

In all of the cases, day or night, the mock poacher heard the UAV well before spotting it or being spotted. Mock poachers reported that being able to hear the UAV helped spot it because the sound provided a general direction to look. The distance between the poacher and the UAV at the time when the poacher heard the UAV is not known because of an inability to record it. To allow the auditory detection of the UAV to be examined, the ability of the poacher to hear the UAV on takeoff was evaluated for all of the quadcopter flights. The quadcopter took approximately 30 seconds after takeoff to ascend to flight altitude and load waypoints before starting its autonomous flight pattern. Based on this knowledge, any mock poacher who heard the UAV within the first 30 seconds could hear the quadcopter at the takeoff location at the 30m-flight altitude. A slight trend between auditory detection and distance was present, however there was no strong evidence of a distance at which the quadcopter is no longer audible. This lack of clear correlation is due, at least in part, to the effects of varying terrain between zones. Figure 9 below shows the locations where the UAV was heard on takeoff for Zone A. The area shaded in red represents the view shed from the UAVs perspective. A ridge running through Zone A blocks line of sight from the takeoff location to the eastern edge of the zone, which is also where the UAV was not heard.

**Figure 9:**
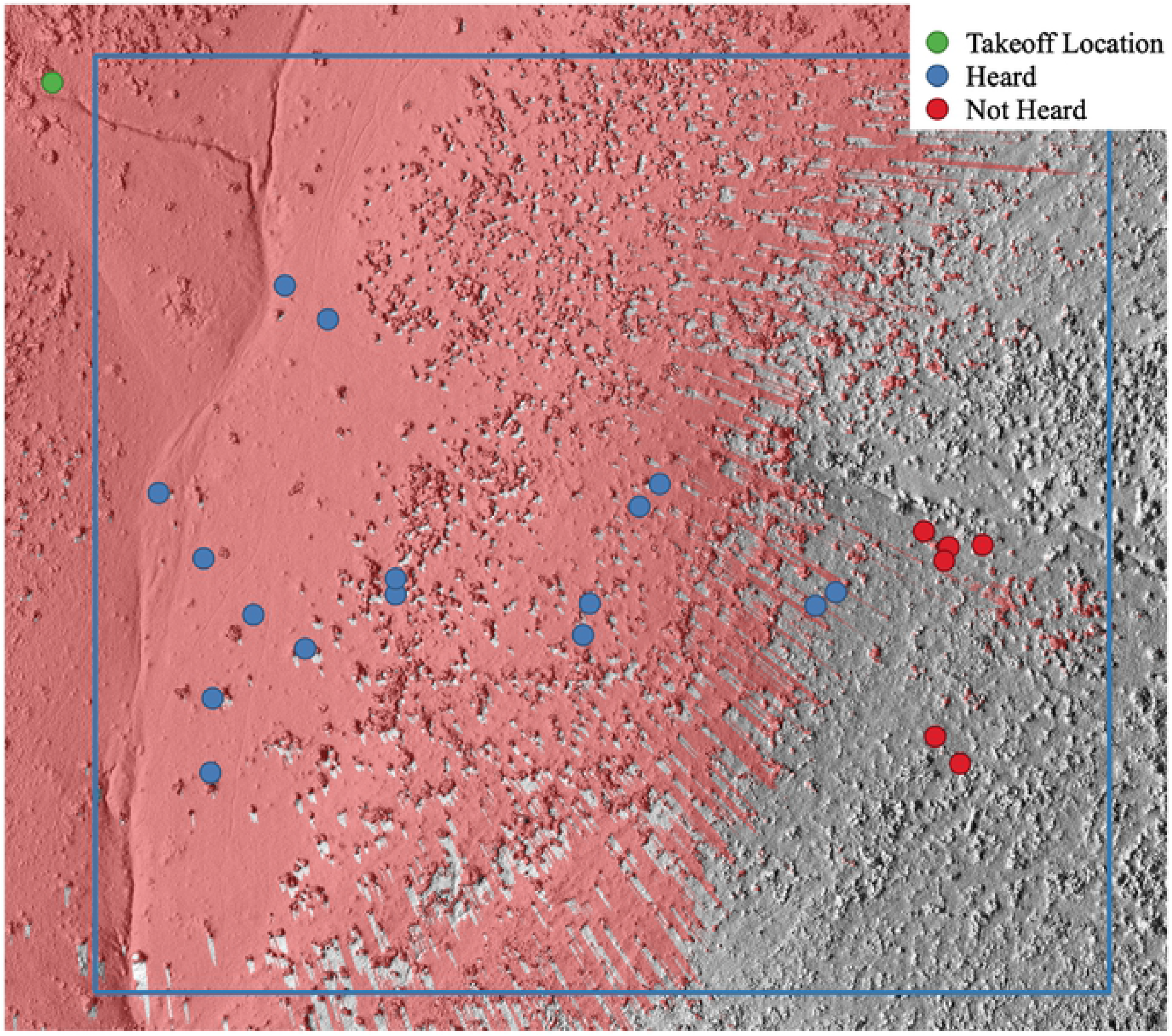
View shed effect on auditory detection

## Summary and Conclusions

In total, 118 trials were completed, providing 236 UAV-poacher interdiction scenarios. Of these, 198 were during the day, 152 with the quadcopter and 46 with the fixed-wing. Live spotting success during the day strongly depended on the hiding behavior of the mock poachers, with the highest success rate being 86% for poachers in the open and the lowest at just 25% for poachers hiding under canopy cover. Both video and still recording were assessed, with the video recording proving to be more effective under the experimental conditions. Below 7.5 m/s the flight speed did not have a significant impact on live spotting success, but observers reported that keeping the flight speed closer to 5 m/s made their work less stressful and more confidently executed. The camera angle also affected confidence of spotting as well as the success rates. A more forward camera angle rather than nadir allowed for the human profile to be better seen as well as remain on the screen longer. For all of the day trials, the mock poachers were easily able to spot the UAV in the sky before being spotted by it.

The remaining 38 interdiction scenarios were at night with the thermal equipped quadcopter. At night the live spotting success rates were much higher due to the thermal camera’s ability to penetrate the bush cover. In the open, 100% of the mock poachers were spotted and the lowest live spotting rate was amongst the bush, with 67% spotted. Although the thermal camera provided an advantage in spotting success, the resolution hindered confidence because of the lack of detail. The mock poachers also struggled with spotting the UAV in the sky due to all of the lights being blacked out and on the majority of the trials the UAV was not spotted.

In contrast to spotting the UAV, the mock poachers had no trouble hearing the UAV from a great distance and used this to help visually search for it. There was no clear correlation between hearing the UAV and distance because the terrain varied greatly between the different study zones and geographic obstructions had a large effect on from how far the UAV could be heard. While the distance at which the UAV could be heard varied greatly, the mock poacher heard the UAV long before the UAV could spot it.

During the day, in every trial, the mock poachers spotted the UAV before being detected. Hiding under cover of canopy after hearing and seeing the UAV approaching reduced the poacher’s chances of being spotted by the UAV to as low as 25%. Much of the mock poacher’s success resulted from the ability to ascertain the direction of the noise source of the UAV. In future studies, that ability could be significantly reduced by flying irregular patterns and/or by employing more than one aerial vehicle. In patrol, the spotting success, whether 25% or higher, would still be a deterrent in countries that enforce stiff penalties resulting from being caught and prosecuted. In interdiction, it could give a poacher time to react and choose to flee the area or hide under cover until the UAV leaves the area. If the poacher were not able to ascertain the direction of the UAV noise source, the poacher might then chose to hide under cover rather than flee. In the interdiction scenario, with a sufficient number of UAVs, the poachers’ movements could be followed and other information about the poachers gathered.

At night, the scenario was similar to during the day, except that, at night the mock poachers struggled to see the UAV in most of the trials. The thermal camera provided an advantage to the UAV by allowing it to better detect humans, achieving a greater than 65% success rate of live spotting across all hiding behaviors. The confusion caused by not being able to pinpoint the location of the UAV created an advantage at night to the security. This uncertainty and the noise that the UAV makes would be ominous to poachers, and therefore a strong deterrent. This makes the UAV a good covert tool, however, it is still better suited as a deterrent because it can still be heard from afar.

For maximum effectiveness across all platforms and times of daytime, the flight speeds should be maximized to give the poachers the least amount of time to react after hearing or seeing the UAV. The limiting factor for flight speed will be the ability of observers to successfully spot poachers without missing people or getting overly fatigued. A forward facing camera angle, preferably around 45° off nadir was found to aid with live spotting success because it increased the time poachers are on screen and it made their profile more visible. Finally, using video allowed for both less interruption as well as more information saved for the post-processing review.

In this study, the quadcopter proved more successful than the fixed-wing, primarily due to its gimbaled camera, higher quality live video transmission, and slower flight speeds. With further improvements the fixed-wing would be expected to provide similar success rates to the quad during the day while also covering more ground and staying aloft longer. At night the fixed-wing would be more difficult to implement due to the need to take off and land on a clearly lit runway.

While the vehicles in this study proved capable of spotting poachers, for them to act as a better covert tool, the noise signature could be reduced. Choosing quieter propulsion systems or flying at higher altitudes would achieve the necessary noise reduction, and higher altitudes would also allow each transect to cover a wider strip of land. However, flying at higher altitudes would also compound the issue of spotting small figures live. To assist in spotting and positively identifying poachers, the use of larger screens and higher quality video for live viewing would help observers be more effective. At night, the slow frame rate and resolution of the thermal camera was a hindrance so a higher resolution thermal camera, with higher frame rate would also be an improvement. During the day, image quality was not the issue; instead human contrast against the background terrain was the issue. In the future, a computer algorithm that highlights possible shapes could help a human observer be more successful and become less fatigued during observation. Despite this need for technical improvements, UAVs were shown, as is, to be an effective tool for deterring poachers because of their ability to spot poachers even with significant auditory presence. Given that the UAVs employed in the study are readily available and a comparatively inexpensive tool both in product cost and in training, the findings support that the integration of UAVs into current anti-poaching patrol and interdiction efforts be aggressively pursued.

## Acknowledgments

The authors would like to acknowledge the field unit consisting of Christina Burnham, Casey Davidson Ciaran Gallagher, Jordan Hatch, Julia Nelson, Abigale Patten, Ana Solberg, Joshua Spann, Cadman Styers, Jaye Sudweeks, Meredith Tooley, Connor Wheatley, and Evan Winterhalter for their efforts in collected and reducing the field data. The Namibian personnel on the ground led by Stuart Munro, and the faculty advisers and the staff in the study-abroad office and the global training initiative office at North Carolina State University are also acknowledged for their wonderful assistance.

## References

1. Air Shepherd. (2018). Retrieved November 28, 2018, from https://airshepherd.org/

2. Ceballos, G., Ehrlich, P. R., Barnosky, A. D., García, A., Pringle, R. M., & Palmer, T. M. (2015). Accelerated modern human–induced species losses: Entering the sixth mass extinction. Science advances, 1(5), e1400253.

3. Douglas-Hamilton, I. (1987). African elephants: Population trends and their causes. Oryx, 21(1), 11–24. doi:10.1017/S0030605300020433

4. Kamminga, J., Ayele, E., Meratnia, N., & Havinga, P. (2018). Poaching Detection Technologies—A Survey. Sensors, 18(5), 1474.

5. Mulero-Pázmány, M., Stolper, R., Van Essen, L. D., Negro, J. J., & Sassen, T. (2014). Remotely piloted aircraft systems as a rhinoceros anti-poaching tool in Africa. PloS one, 9(1), e83873.

6. Western, D., & Vigne, L. (1985). The deteriorating status of African rhinos. Oryx, 19(4), 215–220. doi:10.1017/S0030605300025643

7. WSDOT (Washington State Department of Transportation). (2008). Aviation Emergency Services Aircrew Training Reference Text.

